# Functionalizing silica sol-gel with entrapped plant virus-based immunosorbent nanoparticles

**DOI:** 10.1101/2021.11.12.468100

**Authors:** Matthew J. McNulty, Naomi Hamada, Jesse Delzio, Liber McKee, Somen Nandi, Marjorie L. Longo, Karen A. McDonald

## Abstract

Advancements in understanding and engineering of virus-based nanomaterials (VBNs) for biomedical applications motivate a need to explore the interfaces between VBNs and other biomedically-relevant chemistries and materials. While several strategies have been used to investigate some of these interfaces with promising initial results, including VBN-containing slow-release implants and VBN-activated bioceramic bone scaffolds, there remains a need to establish VBN-immobilized three dimensional materials that exhibit improved stability and diffusion characteristics for biosensing and other analyte-capture applications. Silica sol-gel chemistries have been researched for biomedical applications over several decades and are well understood; various cellular organisms and biomolecules (e.g., bacteria, algae, enzymes) have been immobilized in silica sol-gels to improve viability, activity, and form factor (i.e., ease of use). Here we present the immobilization of an antibody-binding VBN in silica sol-gel by pore confinement. We have shown that the resulting system is sufficiently diffuse to allow antibodies to migrate in and out of the matrix. We also show that the immobilized VBN is capable of antibody binding and elution functionality under different buffer conditions for multiple use cycles. The promising results of the VBN and silica sol-gel interface indicate a general applicability for VBN-based bioseparations and biosensing applications.

## 1. Introduction

Virus-based nanomaterials (VBNs) are being studied for various medical applications as versatile nanomachines that can be manufactured at an industrial scale with high fidelity and low costs^1,2^. Reports in literature have primarily focused on the design of novel VBNs. With sufficient novelty and value of VBNs having been demonstrated in several application areas, there is a need to consider more advanced VBN-based systems to leverage the potential of existing VBNs and move this technological platform towards the clinic and market.

To date, several studies have explored VBNs as structural and/or functional elements within larger system arrangements. Recent examples include: hot-melt extrusion of trivalent vaccine candidates mixed into slow-release PLGA implants^3^; surface conjugation of osteogenic VBN nanofibers to the surface of 3D-printed bioceramic bone scaffolding^4^; electrostatic layer-by-layer assembly of free-standing VBN biofilms^5^; surface adsorption of immunosorbent VBNs onto gold sensor chips^6^; magnetic particle conjugation to immunosorbent VBNs for protein purification^7^.

Silica sol-gel chemistries represent an alluring set of matrices for bioencapsulation and more advanced VBN-based systems. Extensive literature supports favorable silica sol-gel chemistry characteristics of high structural uniformity, stability, pore size tunability, optical properties, and biodegradability for various biomedical applications^8,9^. A range of live cells^10–12^ and enzymes^13–15^ have been studied for bioencapsulation in silica sol-gel matrices to stabilize viability and activity as well as to improve ease of use for the intended application. There have not yet been any such studies using VBNs, with the exception of a study of viral encapsulation focused on extended release of viral vectors for gene therapy^16^.

In this study we present entrapment and utility of plant virus-based immunosorbent nanoparticles (VINs) in silica sol-gel matrices by pore confinement, representing a novel system configuration for VBNs in general. VINs display antibody-binding proteins on their external coat protein surface, and they have been used as simple and bioregenerable reagents for biosensing and therapeutic antibody purification^6,17,18^. This entrapment of VINs represents the first use of plant virus-based VBNs in a silica sol-gel matrix and the first application of VBN technology for utility in an intact silica sol-gel matrix. We demonstrate that the silica sol-gel matrix can immobilize these large biomolecules over a long-duration (~30 days) in an environment that preserves their immunosorbent capture and elution functionalities and is sufficiently diffusive for antibodies to freely enter and leave the matrix. We also show that the silica sol-gel encapsulated VINs can be used to purify antibodies from a complex mixture, in this case, crude *Nicotiana benthamiana* plant extract (representing the production of therapeutic antibodies in plants, formerly known as molecular pharming^19^), to overcome reusability and bioprocessing challenges encountered when using VINs in solution to purify antibodies.

## 2. Materials & Methods

### 2.1 Virion production

Turnip vein clearing virus (TVCV) presenting a coat protein display of the D and E domains of *Staphylococcus aureus* Protein A was used as the plant virus-based immunosorbent nanoparticles. The VINs were produced according to a previously reported study by agroinfiltration of *N. benthamiana* plants using expression vector pICH25892 and processed with a polyethylene glycol (PEG)-based purification scheme^7^. Final yield was approximately 300 mg VIN per kg *N. benthamiana* leaf tissue, as determined by Bradford total soluble protein assay.

Wild-type tobacco mosaic virus (wt-TMV) was produced via mechanical inoculation of ~5-week old *N. benthamiana* plants by lightly sprinkling three leaves per plant with Celite^®^ 545 (Millipore Sigma, Burlington, MA, USA) as an abrasive aid and gently rubbing 100 µL of 0.01 mg/mL wt-TMV in 0.01 M potassium phosphate buffer pH 7.0 per each of the three leaves. The plant leaves were washed with water 20 minutes after inoculation. Leaf tissue was collected after infection symptoms presented ~1 week post-inoculation and frozen at −80 °C for storage.

Extraction of wt-TMV from frozen *N. bethamiana* leaf tissue was performed with a 5:1 (w/v) extraction ratio with 0.1 M potassium phosphate pH 7.0 with 0.1% (v/v) beta-mercaptoethanol using a chilled mortar and pestle. The plant extract was filtered through three-layered cheese cloth, mixed with equal parts chloroform and n-butanol up to 1:1 (v/v) ratio, centrifuged at 8,000 × g and 4 °C for 10 minutes, and the upper aqueous phase layer was collected. PEG-based precipitation was performed by addition of 4% (w/v) PEG 8,000 and 1% (w/v) NaCl, incubation of the mixture for 30-60 minutes at 4 °C, and centrifugation at 8,000 × g and 4 °C for 15 minutes. The pellet was resuspended in 50 mM Tris-HCl pH 7.0 with a glass rod and let to sit at 4 °C for 30-60 minutes. The resuspended solution was centrifuged again at 8,000 × g and 4 °C for 10 minutes. The resulting supernatant was then ultracentrifuged at 50,000 RPM and 4 °C for 90 minutes using a 70.1 Ti rotor (Beckman Coulter, Brea, CA, USA) with 1 mL 15% sucrose cushion per tube. The pellet was resuspended again in 50 mM Tris-HCl pH 7.0 with a glass rod and let to sit at 4 °C overnight. The resuspended solution was added on top of a 10 – 40% (w/v) sucrose gradient and ultracentrifuged at 30,000 RPM and 4 °C for 90 minutes using a SW40 swinging bucket rotor. A 50% sucrose solution was used as a plug to fractionate and collect the wt-TMV containing solution.

Purification of wt-TMV was also performed according to the VIN purification protocol. Final yield was ~ 800 mg wt-TMV per kg *N. benthamiana* leaf tissue, as determined by UV absorbance A_260nm_ spectroscopy measurements.

The fluorescent reporter particle used in this study, a chemical conjugation of the Cyanine 5 fluorophore to the exterior of the wt-TMV coat protein (Cy5-TMV), was generated using purified wt-TMV and a two-step reaction composed of a diazonium coupling and click reaction step according to a method presented by Bruckman and Steinmetz^20^. Spin filters were used to separate the Cy5-TMV from leftover reaction reagents. Fluorophore dye loading is calculated using extinction coefficients at 260 nm of ε_Cy5_ = 250,000 mL/cm/mg ^20^ and ε_TMV_ = 3 mL/cm/mg ^21^.

### 2.3 Silica sol-gel synthesis

The silica sol-gel synthesis was performed according to the method for entrapment of liposomes in silica gel detailed by Zeno and team^22^. In brief, 3.8 mL of tetramethyl orthosilicate (TMOS) was added dropwise to 2.75 mL of 0.002 M HCl in a beaker chilled by an ice bath. This mixture was tip sonicated for 15 minutes, added to a round-bottom flask, and rotary evaporated at 340 mbar reduced pressure and 50 °C for 2-3 minutes before being passed through a 0.22 µm filter to yield the final silica sol. The sol was combined with 3 parts PBS by volume and 1 part of a PBS solution containing either VIN, wt-TMV, or Cy5-TMV at ~0.300 mg/mL concentration (according to total soluble protein measurement for VIN or UV-vis measurement for TMV).

The prepared solution was aliquoted as 40 µL droplets (or 2 µL droplets for the fluorescent microscope imaging) onto parafilm at room temperature for gel bead formation. Post-gelation (~1 hr) beads were submerged in equilibration buffer (PBS pH 7.0) at 4 °C prior to use.

### 2.4 Binding and elution of human immunoglobulin G

The binding and elution of human immunoglobulin G (hIgG) with gel beads was performed following a series of processing steps for batch operation: equilibration, sample loading, impurity wash, and elution. Equilibration consists of a 150 µL PBS buffer bath to submerge each 40 µL gel bead, which was individually contained in a 2 mL tube, that was nutated at 4 °C for a minimum duration of 24 hours and exchanged with fresh PBS buffer a minimum of four times throughout the duration. Previous reports in literature have provided 24 hours of equilibration prior to silica sol-gel use^23^, presumably to ensure stabilization of internal pore charges.

The hIgG samples were prepared as 80 µL of 0.25 mg/mL hIgG in either PBS or 0.22 m filtered *N. benthamiana* extract. The clarified *N. benthamiana* extract was prepared by 3:1 (v/w) chilled mortar and pestle extraction of 6-week-old *N. benthamiana* leaf tissue frozen at −80 °C, filtration through 4-layered cheesecloth, centrifugation at 8,000 × g and 4 °C for 15 minutes, and an additional filtration through a dead-end 0.22 µm syringe filter. For sample loading, all equilibration buffer was removed and the 80 µL of sample was added to submerge the silica sol-gel bead, which was kept at 4 °C nutating for 24 hours.

Impurity wash consisted of a minimum of four buffer exchanges into fresh PBS buffer over a period of no less than 24 hours at 4 °C while nutating.

Elution consisted of PBS buffer removal, addition of 80 µL 0.1 M glycine buffer pH 2.5, incubation for 4 hours at 4 °C while nutating, recovery of the elution solution, and pH neutralization with 0.1 volumes of 0.1 M Tris-HCl pH 9.0.

### 2.5 Protein analysis

Protein analyses using methods of the Bradford assay, SDS-PAGE, western blot, and transmission electron microscopy (TEM) were performed according to previously reported methods^7^. UV-vis measurements were made using a Quartz SUPRASIL^®^ quartz cuvette (Hellma Analytics, Plainview, NY, USA) and a SpectraMax^®^ M4 spectrophotometer (Molecular Devices, San Jose, CA, USA). Fluorescence microscopy was performed with an Eclipse 80i microscope (Nikon, Tokyo, Japan) equipped with a 4x objective. Immediately before imaging, Cy5-TMV loaded beads were transferred from a PBS bath to a slide and excess PBS was removed with a pipette. All images were collected using the same exposure time.

## 3. Results

An illustration of the silica sol-gel functionalization with entrapped plant virus-based immunosorbent nanoparticles and the example use case presented in this study is depicted in Figure 1.

**Figure 1.**
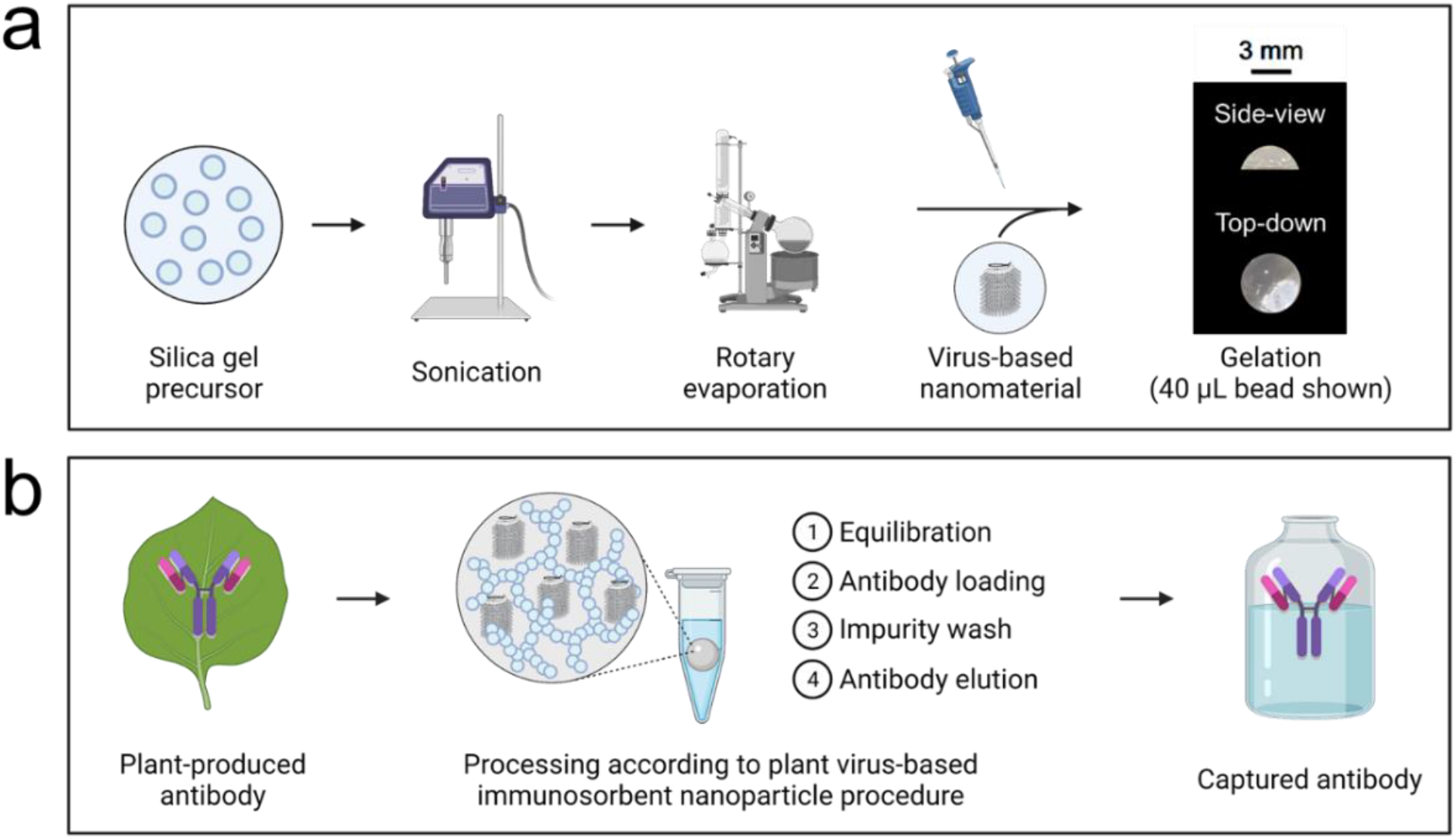
Illustrated schematics of (a) synthesis of sol-gel functionalized with entrapped virus-based nanomaterials (including example photographs of gels synthesized in this study), and (b) the example use case presented in this work of gel-entrapped plant virus-based immunosorbent nanoparticles for affinity purification of plant-made antibodies.

Firstly, a fluorescent reporter VBN was developed to investigate silica sol gel entrapment. As shown in Figure 2a, we produced wt-TMV *in planta* and purified the wt-TMV to ultra-high levels of purity. We then generated Cy5-TMV as a reporter system using previously reported methods to conjugate Cyanine 5 fluorophore to the exterior surface of wt-TMV coat^20^. UV-Vis absorbance spectroscopy 200 – 700 nm spectrums of wt-TMV and Cy5-TMV are comparable to the spectrum results of the previously reported study (Figure 2b). Notably differences of the Cy5-TMV from the wt-TMV include the introduction of an absorbance peak at around 324 nm, which is consistent with the introduction of a diazonium bond (needed for the alkyne addition reaction), and another peak at around 646 nm, which is consistent with the sulfo-Cy5 azide absorbance. We estimate a dye loading of ~32%, which is to say that on average ~680 of an estimated 2,130 coat proteins per assembled virion were conjugated with a Cy5 fluorophore on the exterior surface. Negative stain TEM images were used to confirm structural integrity of the Cy5-TMV (Figure 2c).

**Figure 2.**
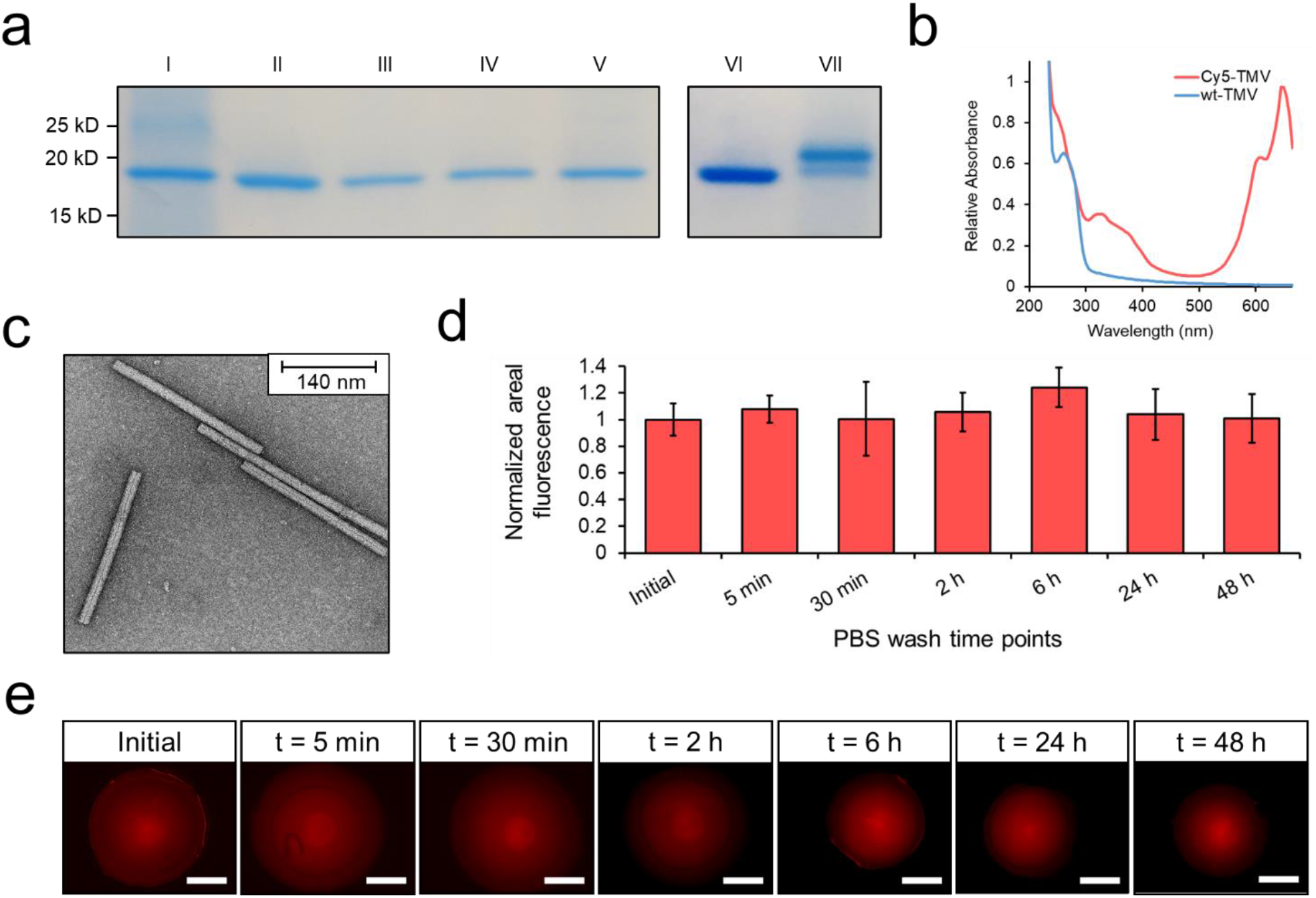
Purification and immobilization of fluorescently labelled plant virions (a) SDS-PAGE results of an ultracentrifuge-based purification of wt-TMV from *Nicotiana benthamiana* leaf tissue, which is then chemically conjugated with Cy5 dye in a two-step reaction to generate Cy5-TMV. Lane definitions: I – filtered plant extract, II – PEG-precipitated pellet resuspension, III – microfuge pellet, IV – ultracentrifuge pellet resuspension, V – ultracentrifuge wash, VI – initial wt-TMV pre-conjugation, VII – Cy5-TMV post-conjugation. (b) UV-Vis absorbance spectra of wt-TMV and Cy5-TMV. (c) Negative stain transmission electron microscope image of the Cy5-TMV. (d) Normalized areal fluorescence per 2 µL volume silica bead containing Cy5-TMV over 48 hours. Beads were exchanged into fresh PBS buffer after each measurement. Error bars represent one standard deviation with biological triplicate. (e) Fluorescent microscope images of the silica sol-gel beads containing Cy5-TMV over 48 hours of submerged gel wash. Scale bars represent 500 µm.

Plant virion entrapment due to crosslinking physical immobilization during silica sol-gel bead synthesis was then assessed using the Cy5-TMV reporter system. Normalized areal fluorescence measurements are shown over the course of a 48-hour study in Figure 2d. Representative fluorescence microscopy images used in the quantitative measurements are shown in Figure 2e. Fluorescence results indicate that the initial Cy5-TMV concentration in the silica sol-gel matrix was maintained throughout the period of examination. There were appreciable decreases in gel bead volume due to minor breakages and shrinkage over the course of multiple microscope images (Supplementary Information, Figure S1). We attribute this observed behavior to the small bead size (2 µL), forceps manipulation, and multiple exposures to a dry environment used for fluorescent microscope imaging. This observation of areal shrinkage was not noted in the beads used for functional testing (40 µL).

There was no evidence of VIN or wt-TMV lost in the PBS wash buffer during standard equilibration to suggest incomplete entrapment. However, we performed an experiment using lower wash buffer volumes (1 vol buffer: 1 vol gel) to improve the limit of detection and found that < 10% of the initial VIN or wt-TMV mass added to the gel was recovered in the wash buffer during equilibration, peaking at an initial brief wash timepoint and decaying rapidly from there (Supplementary Information, Figure S2). This suggests that a small amount of the VIN or wt-TMV was not adequately entrapped during sol-gel synthesis and that it can be readily cleared from solution.

The bind-and-elute immunosorbent functionality of the VIN-entrapped silica sol-gel beads was then examined in this study. Assessment of this functionality using samples of hIgG in PBS and hIgG in clarified *N. benthamiana* plant extract is shown in each condition for two consecutive cycles of bind-and-elute in Figure 3. The results show that the VIN-entrapped beads yielded a high recovery of hIgG in the elution step for both the PBS and plant extract conditions, whereas the TMV-entrapped beads did not.

**Figure 3.**
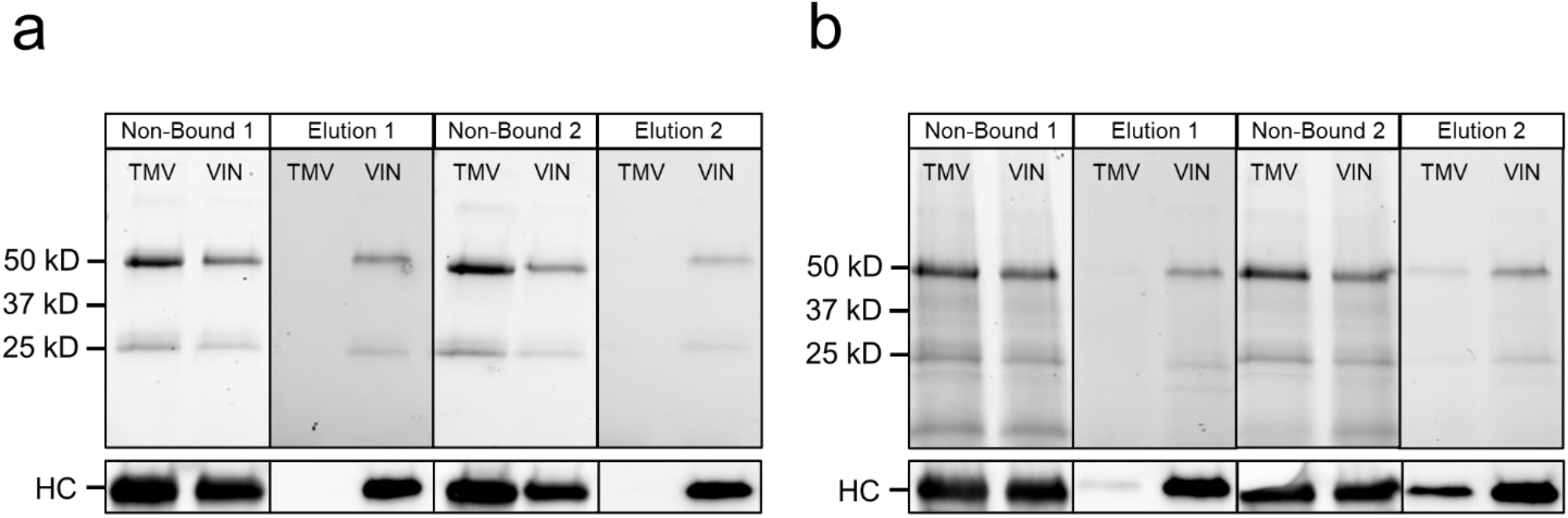
Use of silica sol-gel functionalized with entrapped plant virus-based immunosorbent nanoparticles (VIN) to purify monoclonal antibodies. Reducing condition SDS-PAGE (upper) and western blot (lower) results of the non-bound liquid sample after loading and low pH elution of human immunoglobulin G for a first and second use cycle shown for a sample loading consisting of human immunoglobulin G spiked into a solution of (a) clean PBS and (b) sterile-filtered *Nicotiana benthamiana* extract. Lanes are loaded with fixed volume. HC, heavy chain of the human immunoglobulin G.

The small amount of hIgG in the elution for the wt-TMV negative control using *N. benthamiana* extract (Figure 3b) indicates that the bind-and-elute functionality cannot be solely ascribed to specific Protein A-Fc interactions. Additional negative controls of silica sol-gel synthesized with bovine serum albumin and with no proteinaceous solution also corroborate these results (data not shown). The hIgG recovered in the elution for the plant extract-containing sample condition indicates some degree of non-specific binding interactions which may be present. Minor yellowish coloration of the gel bead, which closely resembled the color of the loaded sample, after impurity washing provides visual support for this observation (data not shown).

The second use cycle of hIgG recovery in elution indicates that the VIN functionality is preserved over multiple uses. A three-week lag period between the gel synthesis and completion of the second use cycle also provides additional support for the entrapment and stability of VIN in the silica sol-gel environment. An increase in hIgG recovery for the second use cycle elution of the wt-TMV bead and plant extract-containing sample condition is observed, which may indicate an undesired behavior for performance such as incomplete hIgG elution from the first use cycle or an additive effect of the non-specific binding mechanism between the first and second use cycle such as what may be observed should a subset of the plant extract constituents accumulatively bind to the silica sol-gel matrices.

The hIgG captured from the plant extract sample was recovered at ~60% purity in both use cycles, based on gel densitometry measurement. The SDS-PAGE results qualitatively support that the hIgG purity was significantly increased from the capture and elution procedure. A reduction in recovery for the second use cycle was observed for hIgG in PBS (~50% first elution) and in plant extract (~80% first elution).

A substantial fraction of the hIgG sample loaded was not bound in either the VIN or wt-TMV conditions. We attribute this to the excess loading concentration, which was used in this proof-of-concept study to determine maximal elution given the employed gel synthesis and operational configuration. Each gel bead was synthesized containing a total of ~3 µg VIN, which, based our previous study of VIN in free solution recovering ~1.5 mg hIgG/mg VIN in excess hIgG conditions, can be estimated to correspond to a binding capacity of ~4.5 µg hIgG. Each gel was incubated in a liquid bath containing ~20 µg hIgG. Based on Bradford total soluble protein assay results, we recovered ~1.9 µg hIgG from PBS for the first cycle elution from the immobilized VIN sol-gel bead, corresponding to ~10% hIgG recovery of the initial sample load. This binding capability represents ~42% of that of VIN in free solution.

## 4. Discussion

In this study, we report encouraging proof-of-concept results for novel VBN entrapment and functionality within the pores of silica sol-gel matrices with broad applicability in protein purification and biosensing. VIN-containing silica sol-gel beads were used to capture hIgG from PBS or plant extract and elute the hIgG in a low pH environment for two consecutive use cycles spanning approximately 30 days post-synthesis. The high surface area to volume ratio of the rod-like VBNs used in this study proved to be an amenable geometry for pore entrapment with considerable active binding site availability without the need for significant optimization.

Future work to investigate the significance of VBN morphology on performance would be valuable for understanding the possible VBN silica gol-gel design space. Relevant characteristics for investigation include the geometry of the virion, either rod-like or icosahedral, the rigidity of the geometry, given that rod-like virions can be classified as stiff (e.g., tobacco mosaic virus) or flexible (e.g., potato virus X), and the size of the virion, which can be readily extended in rod-like virions through genome augmentation.

Non-specific binding interactions of the hIgG to the silica sol-gel were observed in this study, as observed for the wt-TMV and plant extract sample condition. We hypothesize that constituents of the plant extract bind to the silica sol-gel and in turn the extract-gel complex increases the non-specific binding interactions with hIgG. Additionally, we hypothesize that the low pH elution conditions (pH 2.5) and the isoelectric points of the silica sol-gel matrix (pH 2.0 for silica) and hIgG (> pH 6.0) may generate a non-specific binding environment that could result in incomplete hIgG elution into the bulk liquid. This behavior was not observed in the experimental execution of this study but should be considered in the future. The isoelectric points of wt-TMV coat protein, pH ~4^24^, and VIN coat protein fusion, pH ~3.7 as estimated using Expasy (web.expasy.org/compute_pi/), are both net negatively charged at the neutral gelation condition and not expected to non-specifically bind with the silica sol-gel matrix. We do suggest that future works more rigorously resolve concerns of non-specific binding interactions through optimization of silica sol-gel composition, perhaps considering doping (3-aminopropyl) triethoxysilane into the formulation, as has been successfully employed in previous studies^25^.

The reduction of effective binding capacity for silica sol-gel entrapped VIN was within the range of previously reported reductions in enzyme activity; silica sol-gel entrapped horseradish peroxidase and glucose-6-phosphate dehydrogenase enzymes were reported to exhibit specific activity of 73% and 36% of the specific activities of the free enzymes, respectively^23^. Further investigation is required to understand the relative contributions of protein activity modulation, such as from VIN exposure to gelation conditions or the internal pore environment pH, and loss of accessibility of the active site, such as from diffusional limitations in the sol-gel matrix or partitioning of VIN into hIgG-inaccessible silica sol-gel matrix pores.

The findings of this study were consistent with the results of Kangasniemi and team for extended release of adenovirus from silica sol-gel in finding that the VBNs in silica sol-gel were stable for extended durations (weeks to months) and could be evenly distributed in the sol-gel post-entrapment, as noted by the nearly linear VBN release profile during *in vitro* silica dissolution in their study^16^. The fundamental difference in observations was that Kangasniemi and team showed VBN activity to be retained for VBN that were released into free solution via silica dissolution, whereas this study reports VBN activity while entrapped within the silica sol-gel matrices. Our work also presents the first entrapment of plant virus-based VBN in silica sol-gel, which can be used in a wide variety of applications not amenable to mammalian virus-based VBN due to inherent advantages of safety (e.g., non-infectious to humans) and inexpensive and simple production.

The VIN silica sol-gel system presented in this study could serve as the foundation for a robust and reusable platform for biosensing. For example, the VIN system could be readily configured for immunosensing, in which the desired detection antibody would be first introduced to and fixed (via non-covalent bonding) in the VIN-entrapped gel matrix, allowing the subsequent addition of the desired analyte solution for sensing. The detection antibody could be removed via low pH elution and the same VIN-entrapped gel could be reused with a different detection antibody, as desired. This hypothesized example system may provide means for easy reuse and flexibility of sensing targets as well as an enhanced sensitivity over antibody-only systems due to the increased sensing surface area of the structural scaffolding of the VIN. This hypothesis of the potential for increased sensitivity is strongly supported by our recently reported results that VIN coupled to magnetic particles could achieve ~25x higher binding capacity of antibodies as compared to current industry standards for affinity protein capture with magnetic particles on a per unit mass basis^7^.

Future works to further exploit the advantages of the highly tractable silica sol-gel chemistries for more sophisticated geometries and fine-tuned pore architecture would be valuable. For example, a previous study showed synthesis of bimodal pore distribution silica sol-gel in monolithic nanoflow columns in which the smaller pores are used to entrap the ligand to functionalize the column and the larger pores promote bulk flow and diffusion of the target molecule to the ligand active site^26^. This system architecture could be valuable for developing scale-down VBN-based bioseparation or biosensing technologies.

## Abbreviations

Cy5-TMV: Cyanine 5 conjugated TMV
hIgG: human immunoglobulin G
PEG: polyethylene glycol
PLGA: poly(lactic-co-glycolic acid)
TMOS: tetramethyl orthosilicate
TMV: tobacco mosaic virus
TVCV: turnip vein clearing virus.

## 5. Acknowledgements

This material is based upon work supported by NASA under grant or cooperative agreement award number NNX17AJ31G, a NASA Space Technology Research Fellowship (NASA grant number 80NSSC18K1157), the Translational Research Institute through NASA Cooperative Agreement NNX16AO69A, and the National Science Foundation (Grant number DMR – 1806366). Any opinions, findings, and conclusions or recommendations expressed in this material are those of the author(s) and do not necessarily reflect the views of the National Aeronautics and Space Administration (NASA), the Translational Research Institute for Space Health (TRISH), or the National Science Foundation.

The illustrations were created in part using Biorender.com.

