## Supplementary Information for "Functionalizing silica sol-gel with entrapped plant virus-based immunosorbent nanoparticles"

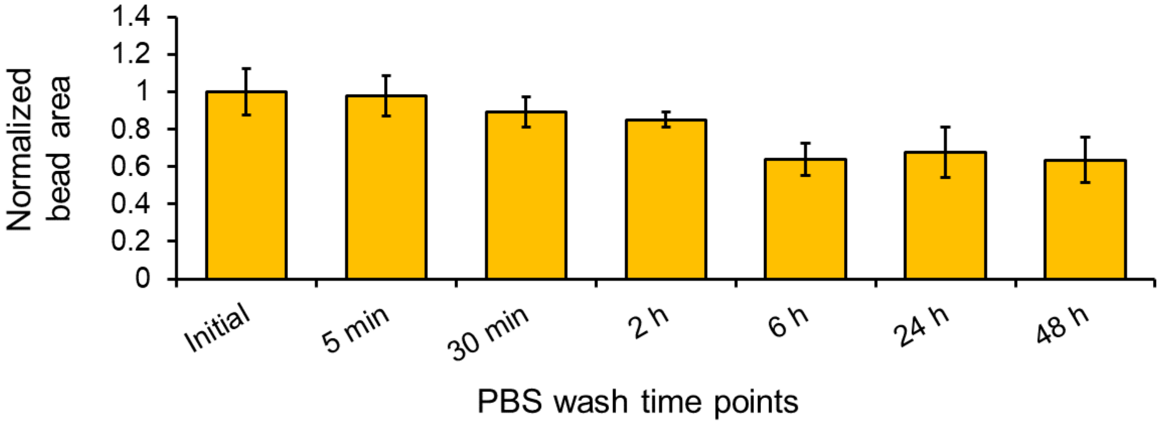


**Figure S1.** Normalized bead area over time for 2 µL volume silica bead containing Cy5-TMV over 48 hours. Beads were exchanged into fresh PBS buffer after each measurement. Error bars represent one standard deviation with biological triplicate.

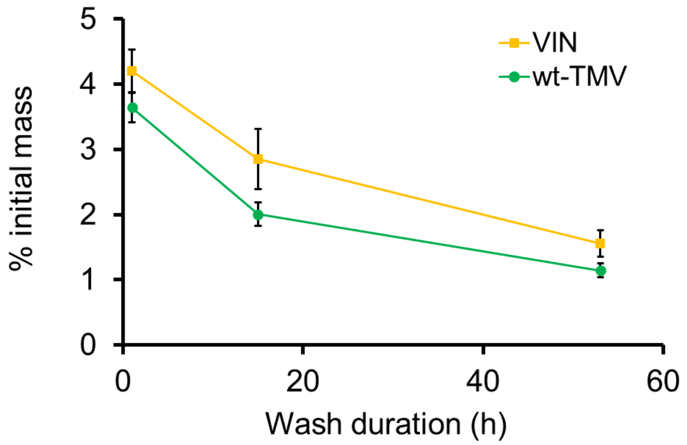


**Figure S2.** UV-vis $A_{280}$ measurements of the PBS wash solution over ~2 days of equilibration reported as a fraction of the initial $A_{280}$ measurement for the VIN or wt-TMV added into the silica sol-gel synthesis. The PBS wash solution was collected at each sample timepoint and exchanged with fresh buffer. This experiment was conducted using 1 volume PBS wash solution per volume silica sol-gel to improve limit of detection. The retention of the PBS wash solution in the buffer exchange (~50% total wash solution) dictates that these results represent an upper bound of the initial mass lost during washing. Error bars represent 1 S.D. using biological triplicate.
